# Alpha-1-antitrypsin and its variant-dependent role in COVID-19 pathogenesis

**DOI:** 10.1101/2020.08.14.248880

**Authors:** Christian S Stevens, Kasopefoluwa Y Oguntuyo, Shreyas Kowdle, Luca Brambilla, Griffin Haas, Aditya Gowlikar, Mohammed NA Siddiquey, Robert M Schilke, Matthew D Woolard, Hongbo Zhang, Joshua A Acklin, Satoshi Ikegame, Chuan-Tien Huang, Jean K Lim, Robert W Cross, Thomas W Geisbert, Stanimir S Ivanov, Jeremy P Kamil, the Alpha-1 Foundation, Benhur Lee

## Abstract

**Rationale:** SARS-CoV-2 entry into host cells is facilitated by endogenous and exogenous proteases that proteolytically activate the spike glycoprotein and antiproteases inhibiting this process. Understanding the key actors in viral entry is crucial for advancing knowledge of virus tropism, pathogenesis, and potential therapeutic targets.

**Objectives:** We aimed to investigate the role of naïve serum and alpha-1-antitrypsin (AAT) in inhibiting protease-mediated SARS-CoV-2 entry and explore the implications of AAT deficiency on susceptibility to different SARS-CoV-2 variants.

**Findings:** Our study demonstrates that naïve serum exhibits significant inhibition of SARS-CoV-2 entry, with AAT identified as the major serum protease inhibitor potently restricting entry. Using pseudoparticles, replication-competent pseudoviruses, and authentic SARS-CoV-2, we show that AAT inhibition occurs at low concentrations compared with those in serum and bronchoalveolar tissues, suggesting physiological relevance. Furthermore, sera from subjects with an AAT-deficient genotype show reduced ability to inhibit entry of both Wuhan-Hu-1 (WT) and B.1.617.2 (Delta) but exhibit no difference in inhibiting B.1.1.529 (Omicron) entry.

**Conclusions:** AAT may have a variant-dependent therapeutic potential against SARS-CoV-2. Our findings highlight the importance of further investigating the complex interplay between proteases, antiproteases, and spike glycoprotein activation in SARS-CoV-2 and other respiratory viruses to identify potential therapeutic targets and improve understanding of disease pathogenesis.

## INTRODUCTION

Severe acute respiratory syndrome coronavirus 2 (SARS-CoV-2), the causative agent of coronavirus disease 2019 (COVID-19), depends on the complex interplay between the virus and host for cellular entry (1–3). Understanding the various steps and factors involved in viral entry is vital to our ability to successfully model the process, identify potential therapeutics, and even predict genetic risk factors. For SARS-CoV-2, this process is mediated by the spike glycoprotein (S). Entry involves not only the simple binding interaction between spike and the entry receptor ACE2, but also the delicate balance of proteases and antiproteases that contribute to proteolytic activation and facilitate viral fusion (4–8). SARS-CoV-2 S undergoes two sequential proteolytic activation steps: first it is cleaved at the S1/S2 polybasic site, and second it is then cleaved at S2′ revealing the fusion peptide (9).

A variety of proteases are implicated in the proteolytic activation of SARS-CoV-2 including furin-like proteases, cathepsins, trypsin, neutrophil elastase, and TMPRSS2 (1, 9–16). This intricate process enables the virus to enter and fuse either at the plasma membrane or within endosomes, engaging different proteases and pathways at each stage. The complexities of viral entry for SARS-CoV-2 are particularly important to understand, as they can be context-dependent, influenced by factors such as local tissue-specific protease and antiprotease milieu, host genotype that impact these milieux, and virus variants that alter susceptibility to cognate protease activation. As an example of the way these relationships can develop and change over time, increasingly efficient proteolytic processing and cell fusion were driving factors in the selective pressure that gave rise to new variants in the first half of the pandemic. New SARS-CoV-2 variants displayed more enhanced cell fusogenicity and proteolytic efficiency relative to earlier strains, starting with the first dominant spike mutation, D614G. Alpha and Beta both showed significantly increased fusogenicity and Delta even more so relative to them (17–21). However, this pattern ended with the rise of the Omicron sublineages as they rely predominantly on endosomal-mediated entry and have significantly reduced fusogenicity (22–26).

Virus-specific mutations are not the only determinant of viral entry and pathogenicity, as various host genotypes can affect virus-host interactions. SNPs in TMPRSS2 and ACE2 for example have been associated with differential COVID-19 risk and/or severity (27–31). Extensive research has been conducted better elucidate the determinants of SARS-CoV-2 pathogenicity. In this study, we investigated the cause of an unexpected inhibition of viral entry by serum samples from patients not previously exposed to SARS-CoV-2. We identify the likely causative factor and discuss its potential role within a complex system where proteases, antiproteases, host genomics, and viral genomics interact and influence each other.

## RESULTS

### Serum inhibition of protease-mediated entry of SARS-CoV-2 pseudoparticles and live virus

SARS-CoV-2 entry is efficiently mediated by a host of endogenous, exogenous, and cell surface proteases. Assays investigating viral entry and entry inhibition should faithfully recapitulate proteolytic activation (Figure 1A) for optimal physiological relevance. Using VSVΔG pseudotyped particles bearing SARS-CoV-2 spike (CoV2pp) we have shown that we can optimize entry efficiency by treatment with exogenous trypsin and followed by treatment with soybean trypsin inhibitor to limit cytotoxicity (32). We tested our trypsin-treated CoV2pp for validity using human serum samples, careful to only using pre-pandemic samples as negative controls (Supplementary Figure 1A-B) (32). As anticipated, sera from SARS-CoV-2 antibody-positive patients exhibit significantly stronger neutralization compared to seronegative sera (Figure 1B-C and Supplementary Figure 1C-F).

**Figure 1.**
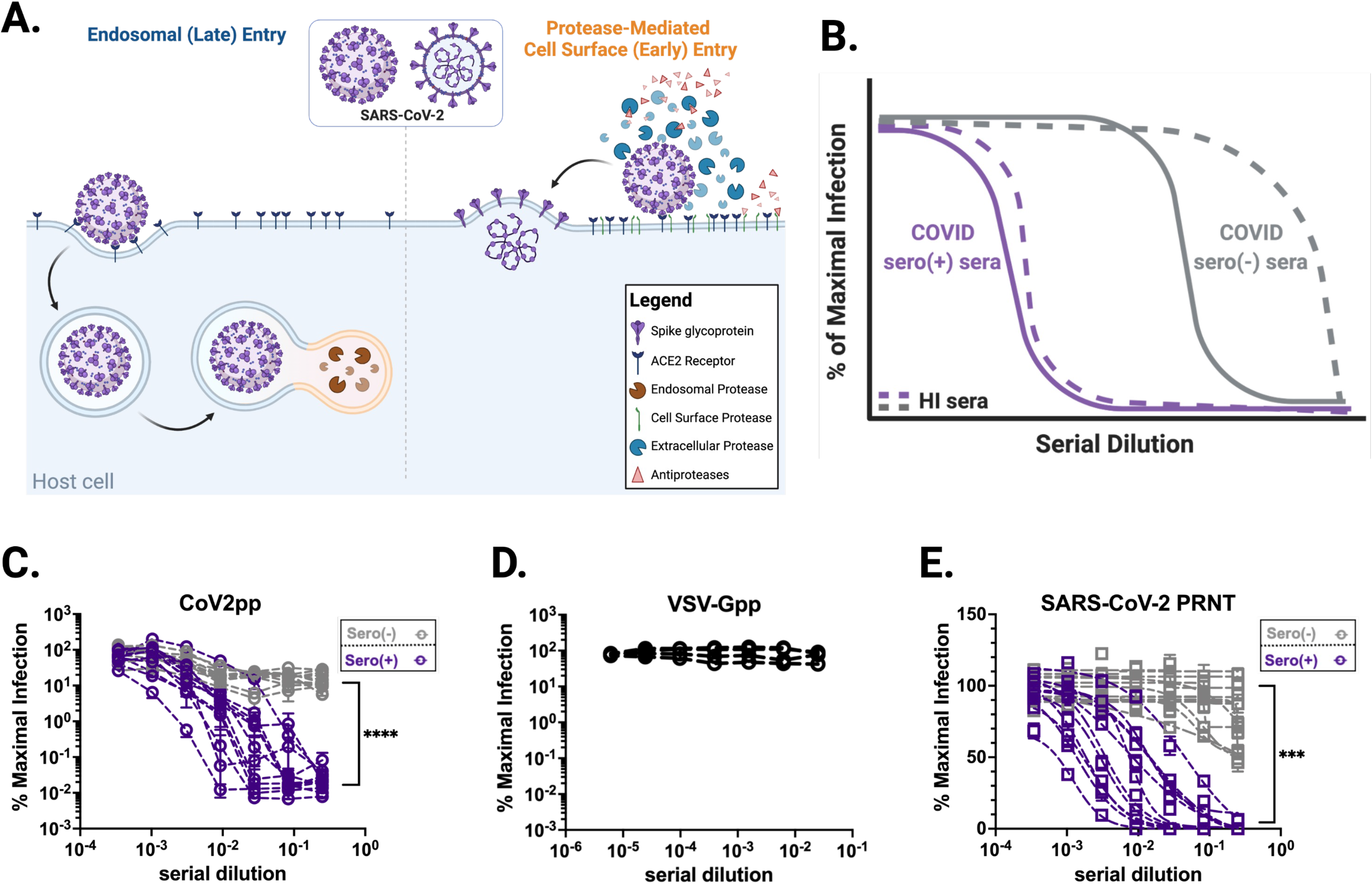
Overview of SARS-CoV-2 entry and inhibition of trypsin treated CoV2pp entry by COVID-19 seronegative sera. **(A)** Overview of SARS-CoV-2 entry. Two modes of entry are displayed: 1) Entry mediated by endosomal proteases such as Cathepsin B/L (late entry) and 2) protease-mediated entry (early entry), driven by cell-surface proteases like TMPRSS2 and extracellular proteases such as trypsin and elastase. Protease-mediated entry may be inhibited by the presence of antiproteases. This model was created in Biorender. **(B)** A representative schematic of entry inhibition of trypsin-treated SARS-CoV-2 pseudotyped particles (CoV2pp) by sera from COVID-19 recovered or naïve individuals (COVID sero (+) and COVID sero (-) sera, respectively). This is a representation of results previously presented in supplemental Figure 3A of Oguntuyo and Stevens et al, mBio 2021, as well as Figure 5B of Nie et al, 2020. Here, sera samples were incubated with trypsin-treated CoV2pp prior to infection of Vero-CCL81 cells. Grey lines represent seronegative sera and purple lines are COVID-19 seropositive sera. The dashed lines are samples that were heat inactivated (HI) prior to use. **(C)** Seronegative and seropositive samples were first identified based on IgG antibodies against Spike (Supplemental Figure 1B). Normalized infection data at the highest and lowest dilutions tested are shown as % maximal infection (media only) with results from seronegative plotted on log scale. Data points represent the mean of neutralizations performed in quadruplicate with SEM bars, each line indicating a sample from a unique donor. Maximal sera inhibition was compared using Welch’s t test. **(D)** SARS-CoV-2 seronegative sera do not inhibit VSV-Gpp. Four serum samples were analyzed each in technical triplicate, means with SEM error bars shown. **(E)** Authentic SARS-CoV-2 is modestly inhibited by seronegative sera. Sera samples also presented in Supplementary Figure 1E were utilized for plaque reduction neutralization experiments (PRNT) with live virus. Presented here are the mean of one experiment done in technical duplicates with SEM error bars. Maximal sera inhibition was compared using Welch’s t test. (ns, not significant; **, p < 0.01; ***, p < 0.005, and ****, p < 0.0001).

Unexpectedly, we found that sera from SARS-CoV-2 naïve patients were also capable of neutralizing these pseudoviruses despite negative Spike ELISA results (Supplementary Figure 1A). An external group at Louisiana State University Health Shreveport (LSUHS) confirmed these findings using the same CoV2pp assay and experimental conditions (Figure 1C and Supplementary Figure 1B, C-F). In both groups, SARS-CoV-2 naïve sera inhibited CoV2pp entry by 90-97% (Supplementary Figure 1E, F). However, seropositive patient sera exhibited inhibition orders of magnitude beyond this threshold, suggesting antibody mediated inhibition of CoV2pp entry (Figure 1C and Supplementary Figure 1C-F). Additionally, using identical serum samples as LSUHS, collaborators at the University of Texas Medical Branch at Galveston (UTMB) also observed significant neutralization of authentic SARS-CoV-2 by seronegative sera, as assayed by a plaque reduction neutralization assay (PRNT) (Fig. 1E and Supplemental Fig. 1G).

During the validation and optimization our CoV2pp system, we described that this inhibition was abolished by heat-inactivation at 56°C for 1hr (32). Here we show a schematic representing entry inhibition by naïve sera from our previously published findings and replicated by other published SARS-CoV-2 neutralization assays using proteolytic activation without sufficient heat inactivation (33–35) (Figure 1B). These findings imply the existence of a heat-labile serum factor or factors capable of inhibiting protease-mediated entry of SARS-CoV-2.

### A serum factor capable of inhibiting protease-mediated entry

Upon observing and verifying our results, we identified alpha-1-antitrypsin (AAT) and alpha-2-macroglobulin (A2M) as abundant and heat labile products in serum that may be responsible for inhibition of protease-mediated entry (36–39). These blood products are typically present in human serum at high concentrations—1.1-2.2 mg/mL for AAT and 2-4 mg/mL for A2M—and have been described to inhibit both exogenous and endogenous proteases (40). A2M and AAT alone are responsible for approximately 10% and 90% of serum antiprotease capacity, respectively (41).

In spite of the name, the primary physiological target of AAT is neutrophil elastase (NE), a protease released by neutrophils and found at high concentrations during acute inflammation, especially in the context of acute respiratory distress syndrome (ARDS) secondary to COVID-19 (42–44). In order to lend weight to the AAT hypothesis, we therefore investigated whether NE, like trypsin or TMPRSS2, was also capable of enhancing CoV2pp entry. We found that NE potently enhances cellular entry relative to untreated particles (Figure 2A), further underlying the potential role of AAT in inhibiting protease-mediate entry *in vivo*.

**Figure 2.**
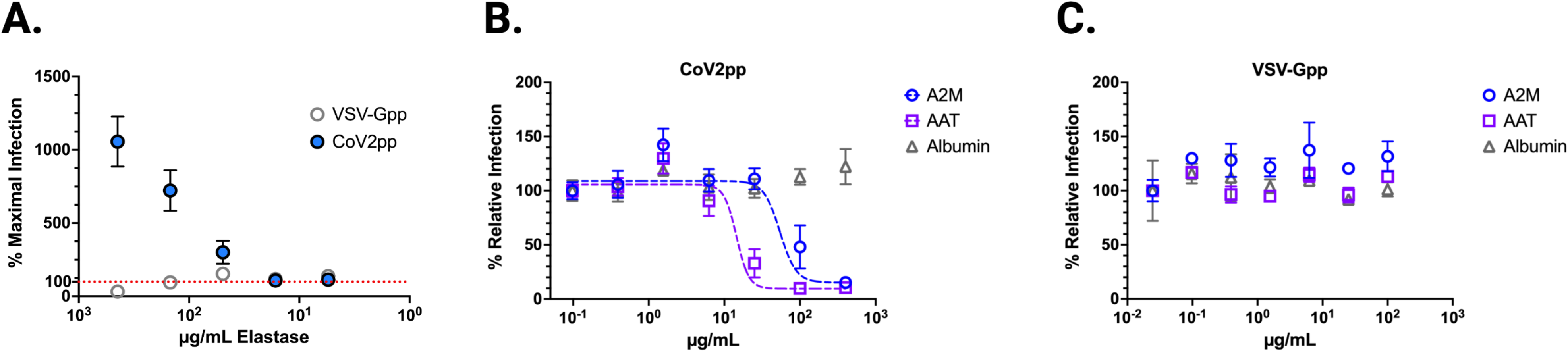
CoV2pp is enhanced by elastase treatment and alpha-1-antitrypsin (AAT) and alpha-2-macroglobulin (A2M) inhibit trypsin-mediated enhancement of CoV2pp entry. **(A)** Treatment of both CoV2pp and VSV-Gpp with elastase in serum-free media. All points are means with SEM bars for samples performed in technical triplicate. Red dotted line marks normalized maximal infection level (100%). **(B)** AAT and A2M inhibit trypsin-mediated enhancement of CoV2pp. Trypsin-treated pseudoparticles were diluted in serum free media, then used to infect Vero-CCL81 cells in the presence of the indicated concentrations of albumin, AAT, or A2M. Data are from two independent experiments and are presented as percent relative infection where each concentration was normalized to the lowest concentration of the test reagent used. **(C)** Performed identical to (B), AAT, A2M, and albumin have no effect on VSV-Gpp entry.

To assess whether AAT and/or A2M alone could inhibit protease-mediated CoV2pp entry, we treated with each at the time of infection and observed dose-dependent entry inhibition by both AAT and A2M, with IC50s of 14.47µg/mL and 54.20µg/mL, respectively (Figure 2B), values that are 50-100-fold below their concentration in serum and bronchoalveolar lavage (BAL) (36, 40, 45). Importantly, neither protein inhibited VSV-Gpp, replicating the inhibitory effect of naïve serum (Figure 2C). Albumin, the most abundant protein in blood, showed no significant reduction of entry of either CoV2pp or VSV-Gpp (Figure 2B), underscoring that the inhibitory effects of AAT and A2M on CoV2-S mediated entry was specific.

While these findings suggest that AAT—and to a lesser extent A2M—can inhibit exogenous proteases known to enhance SARS-CoV-2 entry, SARS-CoV-2 infection is also mediated by proteases at the cell surface as well as endosomal proteases (Figure 1A). TMPRSS2 and cathepsin L are well-characterized examples of cell surface and endosomal proteases, respectively. AAT and A2M are secreted extracellular proteins that can access the former but likely not the latter. Therefore, we sought to investigate whether either protein could inhibit TMPRSS2-mediated SARS-CoV-2 entry. We first showed that saturating amounts of nafamostat mesylate, a specific inhibitor of TMPRSS2, maximally inhibited ∼80% of CoV2pp entry in 293T-ACE2/TMPRSS2 cells (Figure 3A) but had no effect on isogenic 293T-ACE2 cells (Figure 3B). This is consistent with the use of endosomal proteases such as Cathepsin L in the absence of TMPRSS2 or other exogenous proteases. Soluble Spike receptor binding domain (sRBD) completely abolished CoV2pp entry in both cell lines confirming that entry was still entirely ACE2-dependent (Figure 3A-B).

**Figure 3.**
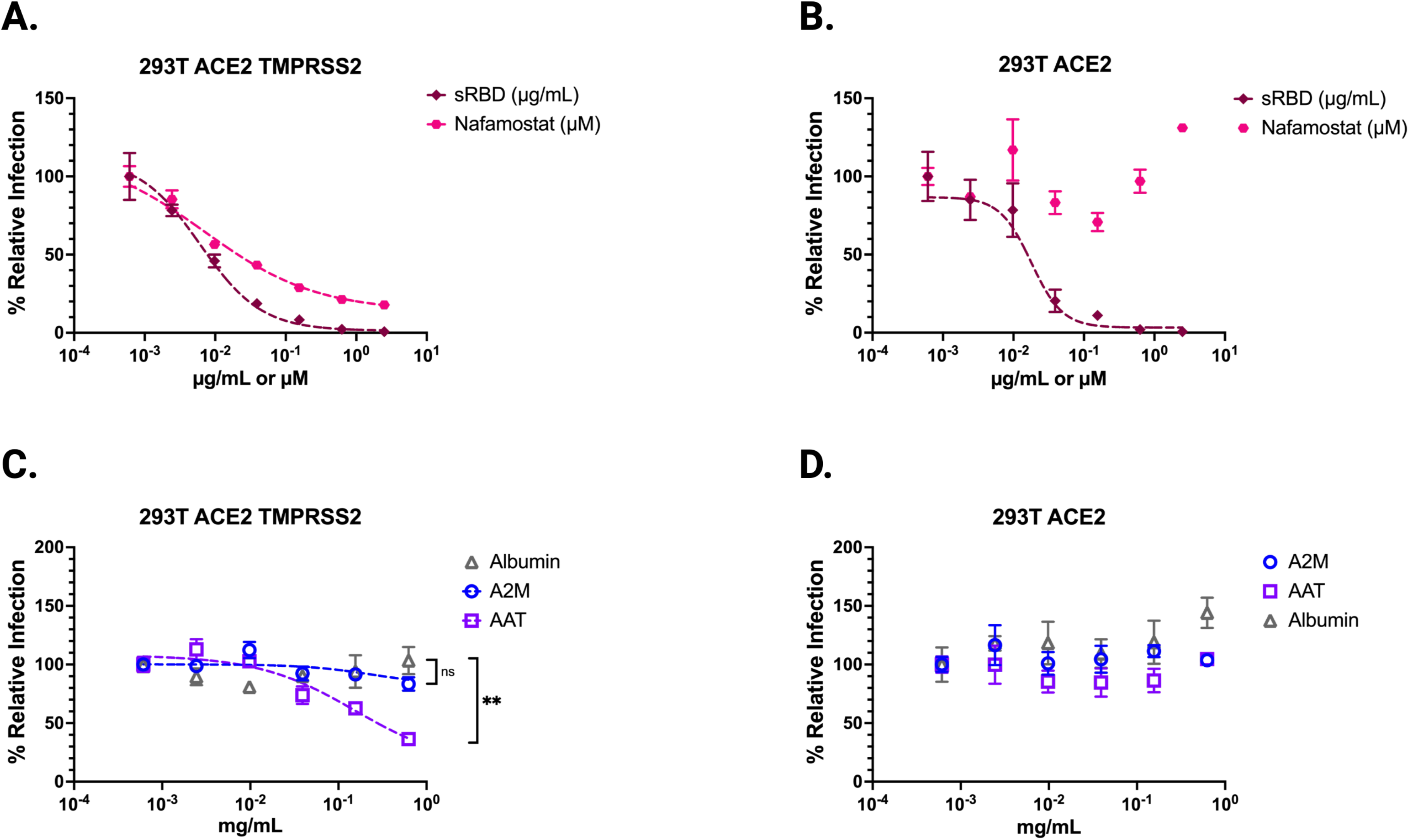
Alpha-1-antitrypsin (AAT) inhibits TMPRSS2 mediated enhancement of CoV2pp entry. **(A,B)** CoV2pp were mixed with a serial dilution of either Nafamostat or sRBD prior to infection of isogenic cells stably expressing (A) ACE2 and TMPRSS2 or **(B)** ACE2. **(C)** AAT inhibits TMPRSS2-mediated enhancement of CoV2pp entry. CoV2pp not treated with trypsin were diluted in DMEM+10% FBS and utilized to infect 293T-ACE2+TMPRSS2 clonal in the presence of the indicated concentrations of A2M, AAT, or Albumin. **(D)** CoV2pp, treated exactly as in **(C)**, were used to infect a 293T-ACE2 clonal cell line. Presented here are the results of an experiment done in technical triplicates. Error bars show SEM and data were fit using variable slope, 4-parameter logistics regression curve (robust fitting method). Significance calculated using a Welch’s T Test on the Area Under the Curve for each condition. (ns, not significant; **, p < 0.01; ***, p < 0.005, and ****, p < 0.0001).

To examine inhibition of TMPRSS2-mediated entry by AAT and A2M, we infected both cell lines with non-protease-treated CoV2pp. We observed that AAT inhibited CoV2pp entry into 293T-ACE/TMPRSS2 cells, but not 293T-ACE2 cells (Figure 3C, D). A2M and albumin both displayed no entry inhibition at the concentrations tested. AAT inhibition of entry into the 293T ACE2-TMPRSS2 cells resulted in approximately a 70% drop in relative infection (Figure 3C). This finding indicates that AAT inhibition accounts for much of the entry attributed to TMPRSS2 enhancement, given that the use of nafamostat mesylate resulted in a maximal inhibition of 80% in the 293T ACE2-TMPRSS2 cells (Figure 3A). This observation suggests that AAT plays a significant role in inhibiting TMPRSS2-mediated entry even at concentrations far below those seen in serum and bronchioalveolar lavage (BAL) (1.1 – 2.2 milligrams/ml) (36, 40, 45).

In summary, our findings provide evidence that AAT, and to a lesser extent A2M, can inhibit protease-mediated entry of SARS-CoV-2 in a cell culture model. AAT demonstrated the ability to inhibit not only exogenous proteases like trypsin but also cell surface protease TMPRSS2, which plays a crucial role in SARS-CoV-2 entry. These results highlight the potential role of these serum factors in modulating viral entry *in vivo*.

### AAT inhibits protease-mediated entry of authentic SARS-CoV-2

To test whether these findings held for authentic SARS-CoV-2, we infected 293T-ACE2 cells and treated them with elastase, trypsin, or a cathepsin inhibitor (E64). We measured the fraction of cells infected with SARS-CoV-2 at 6, 12, 24, and 36 hours post-infection. Consistent with our CoV2pp observations, both elastase and trypsin significantly enhanced entry, while E64 inhibited infection as expected, by inhibiting the cathepsin-mediated pathway utilized when exogenous proteases and TMPRSS2 are absent (Figure 4A).

**Figure 4.**
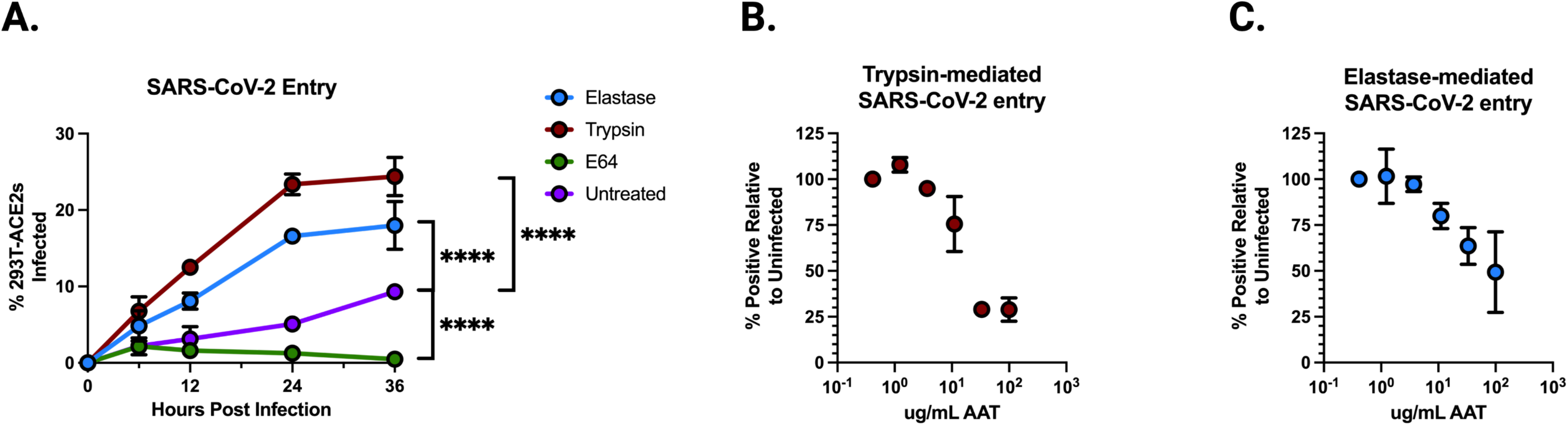
Alpha-1-antitrypsin (AAT) inhibits protease-mediated enhancement of authentic SARS-CoV-2 entry. **(A)** Authentic SARS-CoV-2 entry in 293T-ACE2 clonal cells over 36 hours under treatment of elastase, trypsin, E64, or untreated. Significance calculated using a Welch’s T Test on the Area Under the Curve for each condition. **(B)** Authentic SARS-CoV-2 entry in 293T-ACE2 cells mediated by trypsin, treated by increasing concentrations of AAT, collected at 0, 12, 24, or 36 hours post-infection. **(C)** Authentic SARS-CoV-2 entry in 293T-ACE2 cells mediated by elastase, treated by increasing concentrations of AAT. Data points are means +/-SEM from a representative experiment performed in triplicate. (ns, not significant; **, p < 0.01; ***, p < 0.005, and ****, p < 0.0001).

Next, we performed serial dilutions of AAT in the presence of trypsin- and elastase-treated SARS-CoV-2 at 24 hours post-infection (Figure 4B,C). We observed a dose-dependent inhibition of protease-mediated entry, with potent inhibition at concentrations far below those present in serum and bronchoalveolar lavage fluid. These results are consistent with those obtained using the CoV2pp system, further demonstrating that AAT can inhibit protease-mediated entry of live SARS-CoV-2.

In conclusion, our experiments with authentic SARS-CoV-2 validate the findings obtained using the CoV2pp system, showing that AAT can effectively inhibit protease-mediated entry of the virus. This highlights the potential physiological relevance of AAT in modulating SARS-CoV-2 infection.

### Serum-inhibition of rVSVCov2 entry is reduced in subjects with AAT deficient genotypes

Recognizing AATs potential as an inhibitor of protease-mediated entry enhancement, we aimed to better elucidate the potential physiological relevance of these findings to individuals of various AAT genotypes. AAT levels in serum varies across individuals and are determined by co-dominant alleles in the SERPINA1 gene. The four genotypes investigated here are designated as PI*MM, PI*MS, PI*MZ, and PI*ZZ. Typically, PI*MM individuals have normal AAT levels, while PI*MS and PI*MZ individuals have varying amounts that range from near to sub-normal AAT levels. PI*ZZ individuals are considered AAT deficient (AATD) (Figure 5A).

**Figure 5.**
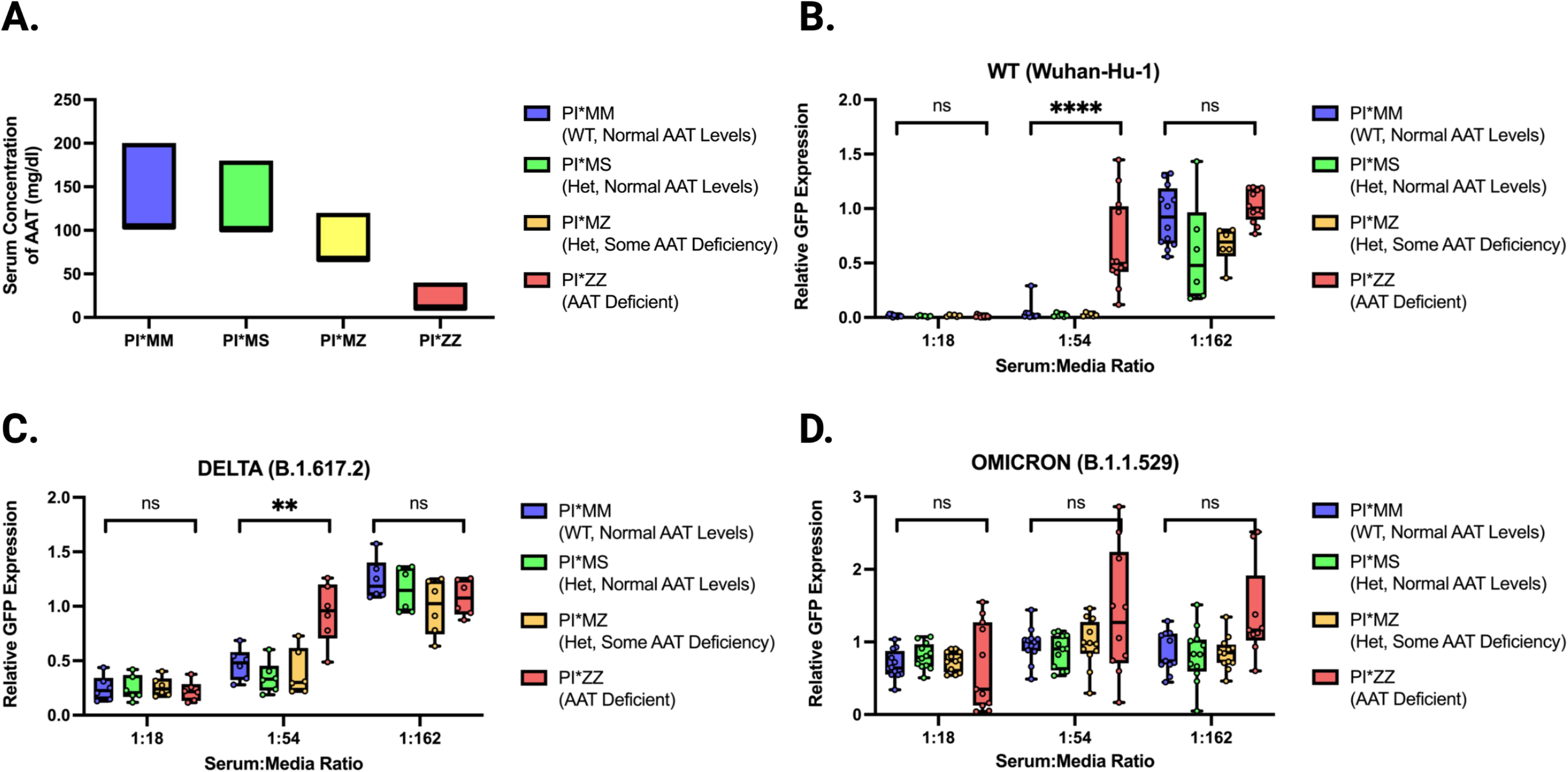
rVSV-CoV2 entry inhibition by serum from subjects with variable AAT genotypes. (A) Expected range of concentrations of AAT in serum relative to AAT genotype (data from Lopes et al (83)). **(B)** WT rVSV-CoV2 entry inhibition by serum from 4 PI*MM subjects, 2 PI*MS subjects, 2 PI*MZ subjects, and 4 PI*ZZ subjects performed in technical triplicate. Subject serum samples are identical for all data shown across all variants. Boxes span 25-75^th^ percentiles and median is noted. Whiskers span minimum to maximum. **(C)** Delta rVSV-CoV2 entry inhibition by serum as described in 3B except only 2 PI*MM and 2 PI*ZZ subjects are shown. **(D)** Omicron rVSV-CoV2 entry inhibition by serum as described in 3B. All data shown performed in technical triplicate. All rVSV-CoV-2 is trypsin-treated. Significance calculated using Welch’s t test (ns, not significant; **, p < 0.01; ***, p < 0.005, and ****, p < 0.0001).

To better elucidate the physiological implications our findings we obtained pre-2020 serum samples from individuals with these four major AAT genotypes and tested their ability to inhibit SARS-CoV-2 Spike-mediated entry. We used replication-competent VSV containing Spike in place of G (rVSV-CoV-2-S) to model SARS-CoV-2 entry. We first infected Vero cells with trypsin-treated WT rVSV-CoV-2-S in the presence of serially diluted serum from the indicated genotypes (Fig. 5B and Supplemental Figure 2A). At 12 hours post-infection, we observed elevated GFP expression at higher serum concentrations for PI*ZZ samples compared to PI*MM, PI*MS, and PI*MZ samples (Figure 5B and Supplemental Figure 2A). This suggests that the most severe AATD genotypes have reduced serum inhibitory potential against protease-mediated entry of WT SARS-CoV-2 Spike, but this reduction is not observed for heterozygotes of the Z and S alleles.

During the SARS-CoV-2 pandemic, several viral lineages have emerged and become dominant. While many similarities exist between all variants of concern (VOCs), some show distinct differences, particularly in viral entry and proteolytic processing of Spike (4, 22, 25). Therefore, we investigated whether the effects of AAT would remain consistent across all SARS-CoV-2 VOCs.

Using our rVSV-CoV-2-S system, we compared entry of isogenic viruses differing only in Spike to model entry by different SARS-CoV-2 VOCs (1). The pattern observed for WT rVSV-CoV-2-S was also seen with Delta rVSV-CoV-2-S containing the Spike protein from B.1.617.2. This VOC became dominant in mid to late 2021, and AATD sera were still less able to inhibit protease-mediated entry (Figure 5C).

Finally, we examined Omicron rVSV-CoV2 containing the Spike protein from B.1.1.529. The Omicron lineage became dominant in 2022 and has remained the dominant lineage since (46), overtaking Delta as the primary VOC in circulation. Notably, Omicron has been shown to undergo less enhancement of entry by proteolytic activation of Spike (22, 25). Using Omicron rVSV-CoV2, we observed no significant difference in inhibitory potential between PI*ZZ and other AAT genotypes (Figure 5D), unlike what we observe for Delta and WT (Figure 5B,C). These data suggest that AAT, and consequently AATD, may have less impact on protease-mediated entry for the Omicron variant compared to WT and Delta variants.

## DISCUSSION

In this study, we identify alpha-1 antitrypsin as the primary inhibitor of protease-mediated entry of SARS-CoV-2 in naïve serum. We demonstrated that AAT effectively inhibits entry mediated by trypsin, elastase, or TMPRSS2. Furthermore, we found that serum from individuals with at least one functional AAT allele (PI*MM, PI*MS, PI*MZ) can inhibit protease-mediated entry by both WT and Delta SARS-CoV-2. Notably, serum from subjects homozygous for AATD alleles (PI*ZZ) exhibited reduced ability to inhibit WT and Delta entry. However, genotype-dependent inhibition of entry was not observed for the Omicron variant, as all serum samples—regardless of subject genotype—displayed similar inhibitory potential. This finding highlights the potential for AAT’s role to vary depending on the entry mechanisms predominantly relied on by the virus.

Approximately 116 million individuals worldwide carry AATD alleles—PI*S or PI*Z—and 3.4 million people are homozygous for AATD—PI*SS, PI*SZ, and PI*ZZ (47). In the early stages of the COVID-19 pandemic, studies reported correlations between mortality rates and the geographic distribution of AAT deficiency (48–50). More recent retrospective analyses indicate that heterozygotes for AATD alleles (e.g. PI*MS, PI*MZ) do not appear to face an increased risk of severe COVID-19 (51). This observation aligns with our findings regarding the inhibitory potential of PI*MM, PI*MS, and PI*MZ sera. Unfortunately, determining a clear increased risk of COVID-19 severity for PI*ZZ patients remains elusive, as large-scale population studies have not yet included a sufficient number of these individuals (52–54). Moreover, disentangling the confounding influence of patients’ awareness of their AATD status presents a challenge. In Germany, survey data from March 2021 revealed that individuals diagnosed with AATD were more concerned with being infected, more likely to restrict their social groups in response, and saw a 65% drop in the percent who had been infected relative to the general population (55).

An additional contributing factor in elucidating the risk associated the alpha-1 antitrypsin deficiency is the emergence of the Omicron lineage of SARS-CoV-2, which is less dependent on the protease-mediated entry pathway for efficient infection (25). As Omicron has become the dominant variant, it has affected the potential risk factors for more severe COVID-19. New variants may alter the role of AAT in disease severity, leading to a different risk profile for individuals with AATD. The COVID-AATD Study from Spain, involving over 2000 patients, showed that both AATD mutations and AAT serum concentration below 116 mg/dl were associated with severe COVID-19 (56). One limitation of this study might be that it was conducted from 1 May 2021 to 1 September 2022, and Omicron (BA.1) overtook Delta as the dominant strain in Spain around 3 January 2022 (57). If AATs antiprotease activity is playing a significant role in their findings of increased risk of COVID-19 severity, separating subjects by estimated date of infection may uncover whether the relationship is lessened after January 2022.

AAT has been predominantly described in the SARS-CoV-2 literature for its role as an acute phase protein in modulating the host immune response (58). AAT plays a critical part in modulating inflammation by inhibiting elastase and other factors (59–62). It is involved in the formation of neutrophil extracellular traps (NETs) in acute pneumonia and can modulate activities resulting in downstream IL-6 inhibition, which is heavily implicated in COVID-19 pathogenicity (41–44, 63). In particular, the IL-6:AAT ratio is highly correlated with COVID-19 severity (64). AAT is also noted for its regulatory role in the coagulation cascade which may have relevance in preventing COVID-19 thromboses (43, 65–67). Previous works have also noted the importance of AAT’s ability to inhibit TMPRSS2 (68). Our study further elucidates the antiprotease role of AAT and suggests that the presence or absence of functional AAT could influence the efficiency of viral entry in other respiratory pathogens.

Patients diagnosed with AATD can receive supplementation through an FDA approved medication: either IV or aerosolized AAT. Four clinical trials for use of AAT in the context of COVID-19 have been registered. One never proceeded beyond recruitment (NCT04385836), one was completed but has not published results (NCT04495101), and one was stopped for futility (NCT04547140). One focused on treating with AAT following a diagnosis of Acute Respiratory Distress Syndrome (ARDS) and saw moderate effects, mostly on modulating inflammation (EudraCT 2020-001391-15) (69). AAT as a treatment for proteolytically activated respiratory viruses shows promise as both an immune modulator as well as a protease-inhibitor and would be well served by clinical trials investigating treatment of a larger patient population as well as treatment in the earlier stages of infection.

Despite the modest results seen in clinical trials of AAT supplementation, the clinical implications of these findings for patients with an AAT deficiency are significant. Typical clinical guidance is that AATD patients are only at increased risk of respiratory infections secondary to chronic obstructive pulmonary disease, not primarily due to the lack of functional AAT except in the case of immune dysregulation (70, 71). Our work suggests this may miss the AATs role as an inhibitor of serine proteases and how that might affect respiratory pathogens reliant on proteolytic activation. For example, not only is TMPRSS2 known to be an important protease for proteolytic activation of the surface glycoprotein hemagglutinin in H1, H3, H7, and H10 influenza A viruses (IAV) (72, 73) but AAT specifically has been shown to inhibit protease-mediated entry of H3N2 IAV and influenza B virus (72). Additionally, multiple studies have implicated AAT in HIV entry as well (62, 74, 75). AAT’s role in viral entry may be specific to the reliance on proteolytic processing at or outside the cell. We show that in the case of SARS-CoV-2, new variants may alter that role, leading to a different risk profile for individuals with AATD. It is crucial to continue investigating the relationship between AAT and new variants to better understand the evolving landscape of risk factors for COVID-19. Furthermore, our findings suggest that it may be worthwhile to invest more into investigating the role of AAT in other respiratory viruses mediated by serine proteases (76, 77). A better understanding of the interplay between AAT and these viruses could help identify potential therapeutic targets, improve patient outcomes, and affect clinical guidance for AATD patients in relation to other respiratory pathogens.

In conclusion, our study highlights the critical role of AAT in inhibiting protease-mediated entry of SARS-CoV-2 and its potential implications for treatment, especially in patients with AAT deficiency. As the COVID-19 pandemic evolves with the emergence of new variants, understanding the role of AAT in the context of these variants is crucial for assessing the shifting risk profiles of individuals with AATD. Our findings also suggest that AAT may play a role in the pathogenesis of other respiratory viruses mediated by serine proteases, opening avenues for future research into the therapeutic potential of AAT in treating these infections. Investigating the role of AAT in other respiratory viruses could lead to the identification of potential therapeutic targets, improved patient outcomes, and updated clinical guidance for AATD patients in relation to other respiratory pathogens. Lastly, further investigation is needed to expand our understanding of AAT’s clinical implications, including the efficacy of AAT supplementation for treating respiratory infections and determining the risk factors for AATD patients in the context of COVID-19 and other respiratory illnesses. By deepening our understanding of AAT’s role in viral pathogenicity and its potential as a therapeutic target, we can better inform clinical practice and contribute to improved public health outcomes.

## METHODS

### Maintenance and generation of isogenic cell lines

Vero-CCL81, BHK-21, Bsr-T7 (78), 293T, 293T ACE2, and 293T ACE2-TMPRSS2 cells were cultured in DMEM with 10% heat inactivated FBS at 37°C in the presence of 5% CO2. Isogenic 293T ACE2, and 293T ACE2-TMPRSS2 cell clones were generated by lentivirus transduction to stably express ACE2 only or ACE2 and TMPRSS2. ACE2 expression was under puromycin selection and TMPRSS2 was under blasticidin selection as previously described (32).

### Production of VSVΔG pseudotyped particles and neutralization studies

Detailed protocols for the production and use of standardized VSVpp (CoV2pp and VSV-Gpp) are given in Oguntuyo and Stevens et al (32). Briefly, 293T producer cells were transfected to express the viral surface glycoprotein of interest, infected with VSVΔG-rLuc-G* reporter virus, then virus supernatant collected and clarified 2 days post infection prior to use. Trypsin-treated CoV2pp were treated as previously described (32). All pseudotyped viruses were aliquoted prior to storage at -80°C and tittered prior to usage for neutralization experiments. Neutralization experiments were performed by diluting the appropriate pseudotyped virus with a 4-fold serial dilution of Albumin (Sigma-Aldrich, A8763), alpha-1-antitrypsin (Sigma-Aldrich, SRP6312), alpha-2-macroglobulin (Sigma-Aldrich, SRP6314) or Nafamostat mesylate (Selleckchem, S1386). SARS-CoV-2 soluble RBD (sRBD) with human IgG-Fc was produced using a recombinant Sendai virus expression platform further described below. All infections were processed for detection of Renilla luciferase activity at 20hrs post-infection, and luminescence was read on the Cytation3 (BioTek).

### SARS-CoV-2 plaque reduction neutralization titration (PRNT) by sera

Neutralization experiments with live virus were performed by incubating sera with 50-100 PFU of SARS-CoV-2, isolate USA-WA1/2020 P4, for one hour at 37°C. All sera were diluted in serum free DMEM. Serial dilutions started at a four-fold dilution and went through seven three-fold serial dilutions. The virus-serum mixture was then used to inoculate Vero E6 cells for one hour at 37°C and 5% CO_2_. Cells were overlaid with EMEM medium (no FBS) and 1.25% Avicel, incubated for 3 days, and plaques were counted after staining with 1% crystal violet in formalin.

### SARS-CoV-2 infection of 293T-ACE2 cells and protease treatment

SARS-CoV-2 isolate USA-WA1/2020 (NR-52281) was provided by the Center for Disease Control and Prevention and obtained through BEI Resources, NIAID, NIH. 1×10^6^ 293T cells stably transduced to express ACE2 were plated in a 6 well dish and infected at an MOI of 0.01 for the time period indicated. Just before infection, virus was treated with alpha-1-antitrypsin (Sigma-Aldrich, SRP6312) at the concentration noted, elastase from human leukocytes (Sigma-Aldrich, E8140) at 0.167 mg/mL, TPCK-treated trypsin (Sigma-Aldrich, T1426-1G) at 50 µg/mL, or E-64 (Sigma-Aldrich, E3132) at 100 µM. Cells were harvested into 4% PFA and were allowed to fix for 30 minutes prior to staining for flow cytometry. Infection was determined with mouse anti-SARS-CoV nucleoprotein antibody, directly conjugated to Alexa Fluor 594. Samples were collected on through an Attune NxT Flow Cytometer and data was analyzed using FlowJo software (v10.6.2). All SARS-CoV-2 work was performed in the CDC and USDA-approved BSL-3 facility at the Icahn School of Medicine at Mount Sinai in accordance with institutional biosafety requirements.

### Production of Replication competent VSVΔG with SARS-CoV-2 Spike Variants and Neutralization Studies

VSV-eGFP was cloned into the pEMC vector containing an optimized T7 promotor and hammerhead ribozyme. Original VSV-eGFP sequence was from pVSV-eGFP; a gift from Dr. John Rose (79). pEMC-VSV-eGFP-CoV2 Spike was cloned as follows: human codon optimized SARS-CoV-2 Spike variants with the 21 amino acid truncation of the cytoplasmic tail were inserted into the VSV-G open reading frame (80) (rVSV-CoV2). The Spike transcriptional unit is flanked by MluI and PacI restriction sites. Expression plasmids containing VSV N, P, M, G, and L open reading frames were each cloned into a pCI vector backbone to allow for efficient virus rescue, generating pCI-VSV-N, pCI-VSV-P, pCI-VSV-M, pCI-VSV-G, and pCI-VSV-L.

rVSV-CoV2 was rescued in 4×10^5^ 293T ACE2-TMPRSS2 or BHK-21 ACE2 cells (32) in each well of a 6-well plate. 2000 ng of pEMC-VSV-EGFP-CoV2 spike, 2500 ng of pCAGGS-T7opt (81), 850 ng of pCI-VSV-N, 400 ng of pCI-VSV-P, 100 ug of pCI-VSV-M, 100 ng of pCI-VSV-G, 100 ng of pCI-VSV-L were mixed with 4 ml of Plus reagent and 6.6 ml of Lipofectamine LTX (Invitrogen). 30 minutes later, the transfection mixture was applied to 293T ACE2-TMPRSS2 cells dropwise. Cells were maintained with medium replacement every day for 4-5 days until GFP positive syncytia appeared. Rescued viruses were amplified in VeroCCL81 TMPRSS2 cells (32), harvested after 6 days, stored at -80C. For titration, 5×10^4^ 293T ACE2-TMPRSS2 cells were seeded onto each well of a 96-well plate and 24 hours later were infected with serially diluted rVSV-CoV2 stock. Virus titer (IU/mL) was calculated 10 hours later by counting GFP positive cells on the Celigo imaging cytometer (Nexcelom).

### Human Sera Samples

All patient sera were acquired after approval by the respective institutional review boards (IRBs) and/or equivalent oversight bodies (Bioethics Committee, Independent Ethics Committee), as follows: (i) the Mount Sinai Hospital IRB (New York, NY, USA), (ii) the Louisiana State University Health Sciences Center—Shreveport (LSUHS, LA, USA), and (iii) the Alpha-1 Foundation (University of Florida, Coral Gables, FL, USA). Samples were deidentified at the source institutions or by the respective principal investigators (PIs) of the IRB-approved protocols for sample collection before analyses performed in this study. All necessary patient/participant consent has been obtained, and the appropriate institutional forms have been archived. Specifically, SERPINA1 genotyped sera samples collected before 2019 were obtained from the Alpha-1 Foundation.

### Enzyme-Linked Immunosorbent Assay

Spike ELISAs for patient sera from the Krammer lab were performed in a clinical setting using the two-step protocol previously published (32, 82). Briefly, this involves screening patient sera (at a 1:50 dilution) with the sRBD; samples determined to be positive were further screened at 5 dilutions for reactivity to the spike ectodomain. Background was subtracted from the OD values, samples were determined to be positive if their ODs were ≥3-fold over that of the negative control, and the AUC was calculated in PRISM. ELISAs performed by the LSUHS group utilized the sRBD with a 1:50 dilution of patient serum to screen all samples, followed by use of the spike ectodomain with patient sera at a 1:100 dilution. Background-subtracted OD values are reported for both sets of ELISAs.

### Production of Soluble SARS-CoV-2 Spike Receptor Binding Domain (sRBD)

Sendai virus (SeV) Z strain (AB855655.1) was cloned into a pRS vector backbone with an additional eGFP transcriptional unit upstream of N. The F transcriptional unit was derived from the SeV Fushimi strain (KY295909.1). We then generated an additional transcriptional unit between the P gene and M gene. SARS-CoV-2 Spike receptor binding domain (sRBD), amino acids 319-541, was taken from human codon optimized Spike (MN908947) in a pCAGGS backbone, a gift from Dr. Florian Krammer (82), and was fused to human IgG1 Fc (amino acids 220-449 of P0DOX5.2) at the C-terminus (SeV-Z-eGFP-sRBD).

2×10^5^ Bsr-T7 cells, stably expressing T7-polymerase, were seeded in a 6-well plate. 24 hours later 4 µg of pRS-SeV-Z-eGFP-sRBD, 4 µg of pCAGGS-T7opt, 1.44 µg of SeV-N, 0.77 µg of SeV-P, 0.07 µg of SeV-L were mixed with 5.5 µl of Plus reagent and 8.9 µl of Lipofectamine LTX (Invitrogen). 30 minutes later, the transfection mixture was applied to Bsr-T7 cells dropwise. Medium was replaced with DMEM + 0.2 µg /ml of TPCK-trypsin (Millipore Sigma, #T1426) at one day post transfection, followed by media replacement every day until infection reached confluency. Supernatant was stored at -80C.

Amplification was performed by seeding 5×10^6^ cells in a T175cm^2^-flask one day before infection. Cells were infected by SeV-Z-eGFP-sRBD at an MOI of 0.01 for one hour, followed by a media replacement with 0.2 mg/ml of TPCK-trypsin-containing DMEM. Cells were maintained with medium replacement by the same every day until infection reached confluency. At maximal infection the medium was changed and replaced with plain DMEM. Cells were incubated for additional 24 hours to allow for maximum protein production. Supernatant was collected and centrifuged at 360 g for 5 minutes, then filtered with 0.1 µm filter (Corning 500 mL Vacuum Filter/Storage Bottle System, 0.1 µm Pore). The flow-through was then applied to a Protein G Sepharose (Millipore Sigma, #GE17-0618-01) containing column (5ml polypropylene columns; ThermoFisher, #29922), followed by wash and elution.

### Statistics and reproducibility

All statistical tests were performed using GraphPad Prism 9 software (La Jolla, CA). For all figures, error bars represent standard deviation of the mean. Sample size and replicates for each experiment are indicated in the figure legends. Technical replicates were prepared in parallel within one experiment, and experimental replicates were performed on separate days. Statistical comparisons as noted in figure legends.

## ACKNOWLEDGMENTS

The authors acknowledge the following funding: KYO and CS were supported by Viral-Host Pathogenesis Training Grant T32 AI07647; KYO was additionally supported by F31 AI154739. BL acknowledges flexible funding support from NIH grants R01 AI123449, R21 AI1498033, and the Department of Microbiology and the Ward-Coleman estate for endowing the Ward-Coleman Chairs at the ISMMS. JPK and SSI acknowledge funding from a LSUHS COVID-19 intramural grant. JPK and SSI acknowledge additional funding from NIH grants AI116851 and AI143839, respectively. Figures created with BioRender.com. We thank Randy A. Albrecht for oversight of the conventional BSL3 biocontainment facility. We would also like to acknowledge the Alpha-1 Foundation and the University of Florida which kindly provided serum samples from SERPINA1-genotyped patients. BL wishes to dedicate this paper to Ernest L Robles-Levroney, the first graduate student BL had the privilege to train. Ernie Robles-Levroney was dedicated teacher, role model and trailblazer who passed away unexpectedly during the course of writing this manuscript.

## FIGURES

**Supplemental Figure 1. Spike ELISA data and neutralization curves.** Spike ectodomain ELISAs for **(A)** ISMMS or **(B)** LSUHS samples. Four seronegative and seropositive samples were utilized. Shown are the OD490 values from the 1:100 sera dilution with the median and interquartile range. **(C)** Seronegative and seropositive samples were first identified based on IgG antibodies against Spike (Supplemental Figure 1A). Normalized infection data at the highest and lowest dilutions tested are shown as % maximal infection (media only) with results from seronegative plotted on log scale. Data points represent the mean of neutralizations performed in quadruplicate with SEM bars, each line indicating a sample from a unique donor. Maximal sera inhibition was compared using Welch’s t test. **(D)** Sera inhibition comparison against VSV-Gpp for 4 sera samples collected at ISMMS. Sera inhibition comparisons for **(E)** ISMMS or **(F)** LSUHS. Inhibition of trypsin treated CoV2pp entry by SARS-CoV-2 seropositive and seronegative sera independently observed. Collaborators in a different state independently performed the identical experiment described in Fig. 1A with their own cohort of seropositive and seronegative samples. Data shown are means from technical quadruplicates/sample/dilution. Experiment performed and presented as in Fig. 1C. **(G**) Live SARS-CoV-2 is inhibited by seropositive and seronegative sera. Sera samples presented in Fig. 1C and (F) above were utilized for plaque reduction neutralization experiments (PRNT) with live virus as described in the materials and methods. Presented here are the mean of one experiment done in technical duplicates and error bars show SEM. (ns, not significant; **, p < 0.01; ***, p < 0.005, and ****, p < 0.0001).

**Supplemental Figure 2. Individual curves of rVSV-CoV2 entry inhibition by serum from subjects with variable AAT genotypes in spike variants: (A)** WT (Wuhan-Hu-1) and **(B)** Omicron (B.1.1.529). Error bars show SEM and data were fit using variable slope, 4-parameter logistics regression curve (robust fitting method). **(C)** Estimation plots for area under the curve shown for PI*MM compared to PI*ZZ in WT rVSV-CoV2 and **(D)** PI*MM compared to PI*ZZ in Omicron rVSV-CoV2. Significance calculated using a Welch’s t test on the Area Under the Curve for each condition. (ns, not significant; **, p < 0.01; ***, p < 0.005, and ****, p < 0.0001).

