## Supplemental Figures for "Alpha-1-antitrypsin and its variant-dependent role in COVID-19 pathogenesis"

Supplementary Figures

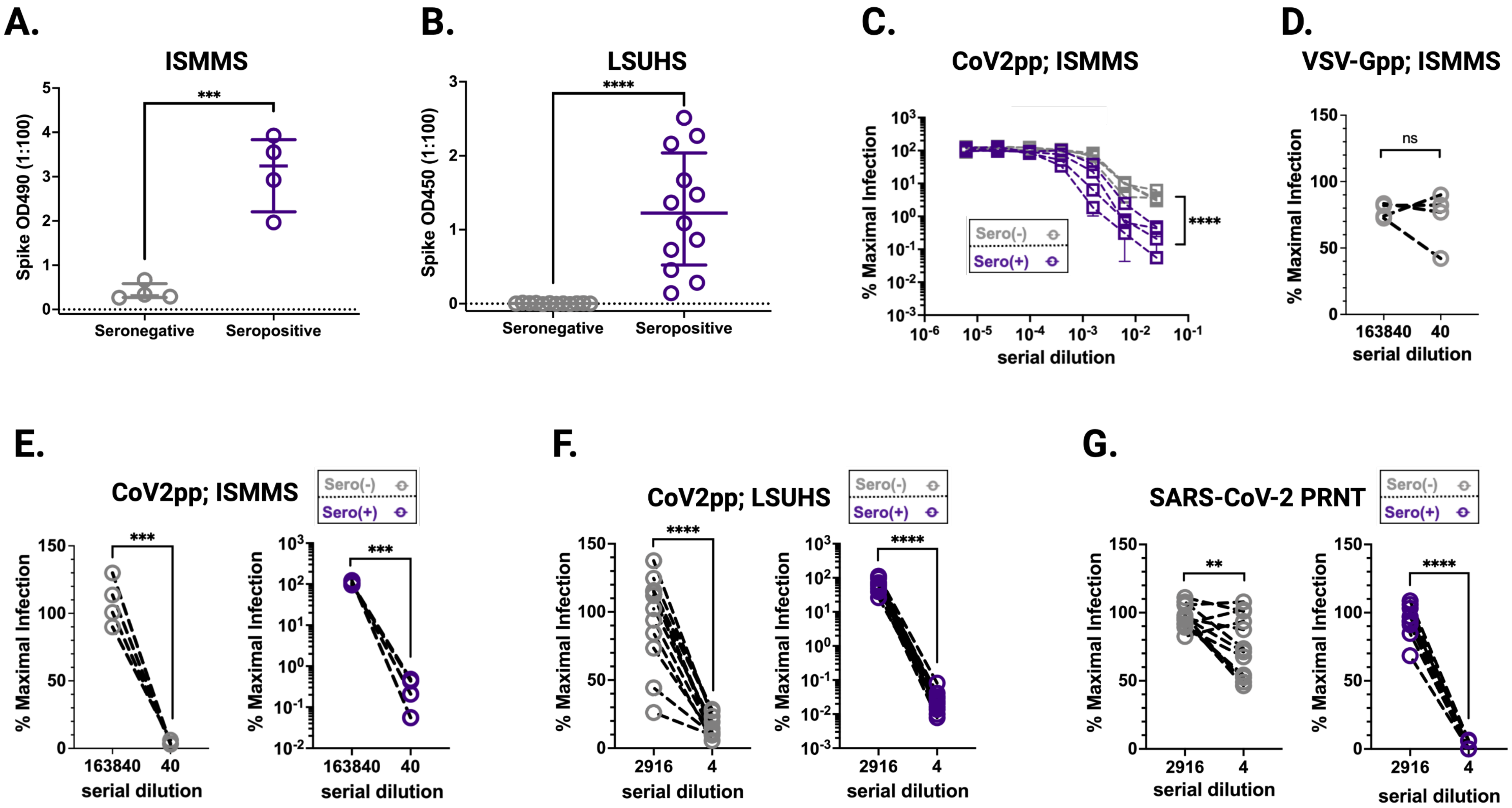

A.
ISMMS
Spike OD490 (1:100)
***
Seronegative
Seropositive
B.
LSUHS
Spike OD450 (1:100)
****
Seronegative
Seropositive
C.
CoV2pp; ISMMS
% Maximal Infection
Sero(-)
Sero(+)
****
serial dilution
D.
VSV-Gpp; ISMMS
% Maximal Infection
ns
163840
40
serial dilution
E.
CoV2pp; ISMMS
Sero(-)
Sero(+)
% Maximal Infection
***
163840
40
serial dilution
F.
CoV2pp; LSUHS
Sero(-)
Sero(+)
% Maximal Infection
****
2916
4
serial dilution
G.
SARS-CoV-2 PRNT
Sero(-)
Sero(+)
% Maximal Infection
**
****
2916
4
serial dilution

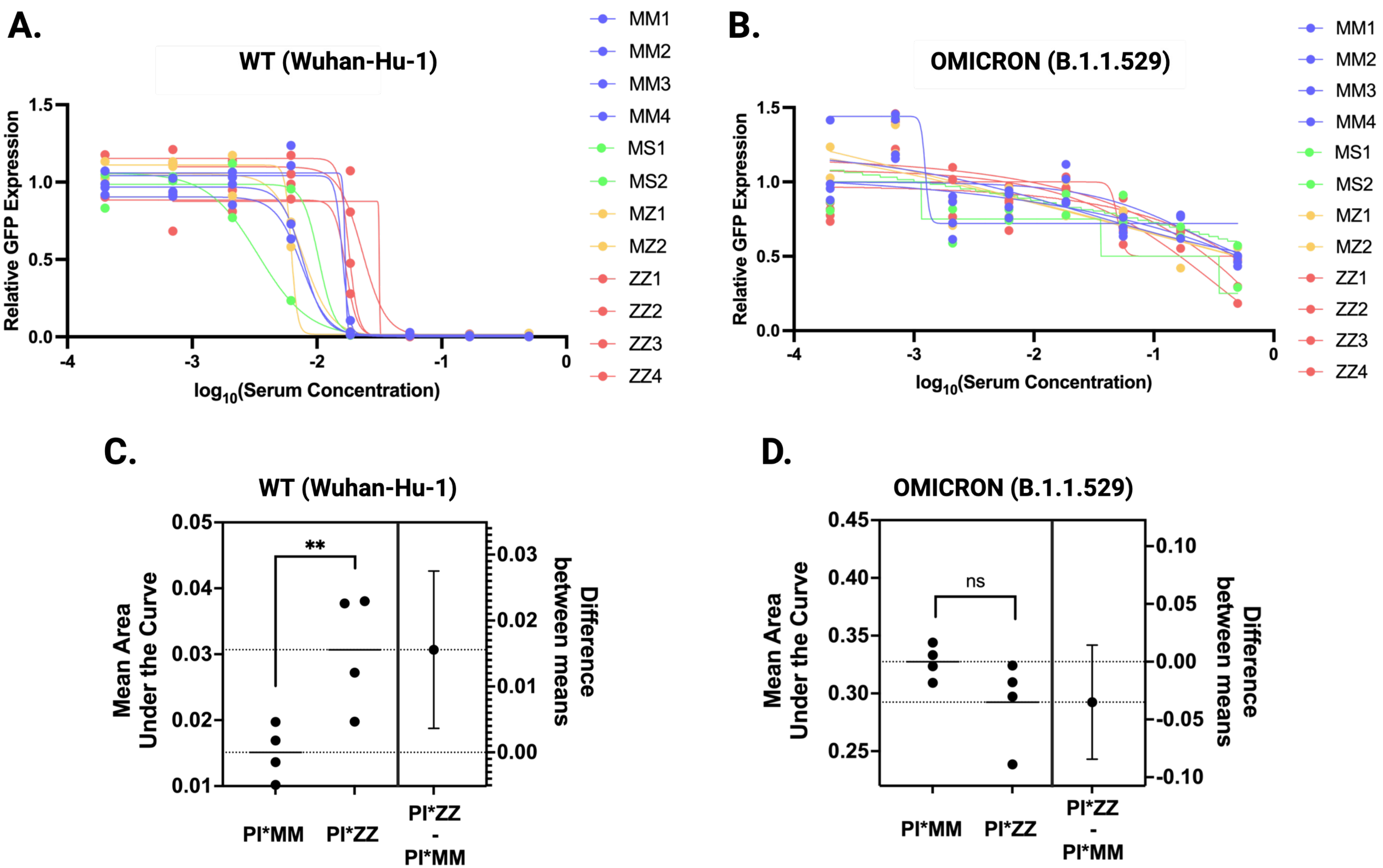
A.
WT (Wuhan-Hu-1)
Relative GFP Expression
log10(Serum Concentration)
MM1
MM2
MM3
MM4
MS1
MS2
MZ1
MZ2
ZZ1
ZZ2
ZZ3
ZZ4
B.
OMICRON (B.1.1.529)
Relative GFP Expression
log10(Serum Concentration)
MM1
MM2
MM3
MM4
MS1
MS2
MZ1
MZ2
ZZ1
ZZ2
ZZ3
ZZ4
C.
WT (Wuhan-Hu-1)
Mean Area Under the Curve
Difference between means
**
PI*MM
PI*ZZ
PI*ZZ - PI*MM
D.
OMICRON (B.1.1.529)
Mean Area Under the Curve
Difference between means
ns
PI*MM
PI*ZZ
PI*ZZ - PI*MM
